# ProNA3D: Distance-Based Analysis of Nucleic Acid-Containing Interfaces

**DOI:** 10.64898/2026.04.16.719043

**Authors:** Luca R. Genz, Maya Topf

**Affiliations:** Research Department of Integrative Virology, Leibniz-Institut für Virologie (LIV); Department of Chemistry, Universität Hamburg; Centre for Structural Systems Biology (CSSB), Hamburg, Germany; Institute for Molecular Virology and Tumorvirology, Universitätsklinikum Hamburg Eppendorf (UKE)

**Keywords:** Protein-nucleic acid interactions, Interface analysis, Structure prediction analysis, RNA secondary structure, CryoEM

## Abstract

Biomolecular interactions are central to many essential cellular processes, but RNA-containing complexes remain challenging to resolve structurally, even as experimental methods and AI-based prediction have expanded structural coverage. Tools for the integrated analysis of complex interfaces remain limited. We present ProNA3D, a tool that provides a unified platform for analyzing protein-nucleic acid and nucleic acid-only complexes, bridging the gap between structure prediction and functional interpretation. ProNA3D supports both experimental and computationally predicted structures, incorporating scoring metrics for AlphaFold3 predictions. It also offers interactive two-dimensional interface visualization and secondary-structure topology plots for RNA and DNA. An interface-based density zoning feature facilitates structure analysis in cryo-EM maps, allowing evaluation of dynamic complexes in the context of heterogeneous density. We demonstrate ProNA3D on diverse complexes solved by X-ray crystallography or cryo-EM, as well as on computational models. For example, in a trimeric complex of HIV-1 RNA and a human antibody, ProNA3D identified a high-connectivity nucleotide with potential functional relevance. Applying ProNA3D to the entire Protein Data Bank revealed distinct interface connectivity trends and interaction modes characteristic of specific complexes (e.g., in methyltransferase-DNA and CRISPR-associated) in nucleic acid-containing interfaces. The method is available as both a UCSF ChimeraX plug-in for visualization and a command-line tool at https://gitlab.com/topf-lab/ProNA3D. In addition, the repository contains the results of the large-scale analyses.

## 2. Introduction

The interaction of proteins with a variety of nucleic acids forms the foundation of many processes within cells, specifically in gene expression. These complexes and contacts are essential not only for mechanisms involving DNA, such as replication, repair, and transcription, but also for the wide spectrum RNA-centered processes [1–4]. Whereas DNA-binding proteins are often associated with transcriptional control [5], RNA-binding proteins (RBPs) participate in a far more extensive set of activities that collectively shape both the transcriptome and the proteome [4,6,7]. RBPs contribute to RNA maturation and processing, including splicing, editing, modification, transport, and translation, acting across cellular compartments from the nucleus to the cytoplasm and organelles [4]. Understanding these cellular functions and the underlying processes depends on the structural analysis of nucleic acids and their interactions with proteins [8,9].

However, experimental structure determination of protein-nucleic acid complexes, particularly those involving RNA, remains challenging. Although techniques such as X-ray crystallography and cryo-electron microscopy (cryo-EM) have advanced structural biology considerably, RNA presents a distinct set of challenges due to its greater flexibility and conformational diversity compared to proteins [10,11]. As a consequence, a significantly lower number of nucleic acid complexes has been experimentally resolved and deposited in the Protein Data Bank (PDB) compared to protein-protein complexes [12]. Recent advances in the field of AI-guided protein-nucleic acid complex structure prediction, such as RoseTTAFoldNA [13] and AlphaFold3 [14], are beginning to address this gap [15]. Although the performance for these complex predictions remains lower than for protein-only assemblies [16–18], these approaches offer a promising opportunity to expand structural coverage and allow the computational analysis of complexes that were previously inaccessible.

A range of computational resources has been developed to support the analysis of protein-nucleic acid and nucleic acid-only complexes (Suppl. Table 1). Tools such as DNAproDB [19] and RNAproDB [20] provide curated topology plots and interactive visualizations of these interactions in three-dimensional space. Secondary structure determination and nucleotide mapping are enabled by tools including DSSR [21], RNAscape [22], and ViennaRNA [23]. Broader interface-focused platforms, including PPI3D [24], PLIP [25], and VoroContacts [26], enable more general interface characterization and analysis. Methods tailored to nucleic acids, such as RNAView [27], further assist in identifying and classifying RNA and DNA base-pairing patterns. Despite their utility, most current platforms are optimized for experimentally resolved structures, and their ability to analyze predicted complex models, which are becoming increasingly important with the advances in AI-based structure prediction, remains limited (Suppl. Table 1). Furthermore, tools for analyzing proteins, RNA, and DNA were often developed independently, resulting in a lack of unified workflows for mixed complexes [28]. In addition, many of these tools are either web servers, restricting users to predefined functionalities, or available only as command-line packages without integrated, interactive visualization. Consequently, there is a need for software that can be used directly within an accessible graphical interface, supports both experimental and computationally predicted complexes, and allows for broad-complex-range (nucleic acid and protein) interface detection and analysis.

Here, we introduce ProNA3D, a UCSF ChimeraX [29] plug-in developed to provide an integrated and accessible environment for the analysis of protein-nucleic acid, and nucleic-acid-only complexes (Fig. 1, Suppl. Table 1). Building on our previously published PICKLUSTER plug-in [30] for protein complex interface analysis, ProNA3D extends interface detection and the identification of spatially separated “sub-interfaces” to a broader range of biomolecular assemblies. It supports the analysis of both experimentally determined structures and computationally predicted models, including scoring metrics for AlphaFold3 predictions. Additionally, ProNA3D provides an interactive and flexible two-dimensional interface visualization and a secondary-structure topology viewer for RNA and DNA complexes, with features for the examination of protein contacts. Finally, ProNA3D enables assessment in the context of cryo-EM maps through density zoning, helping interrogate interfaces and flexible regions that commonly lie in low-resolution areas, especially in composite maps and RNA-containing complexes. Together, these features provide a unified approach for the analysis of protein and nucleic-acid interactions.

**Figure 1:**
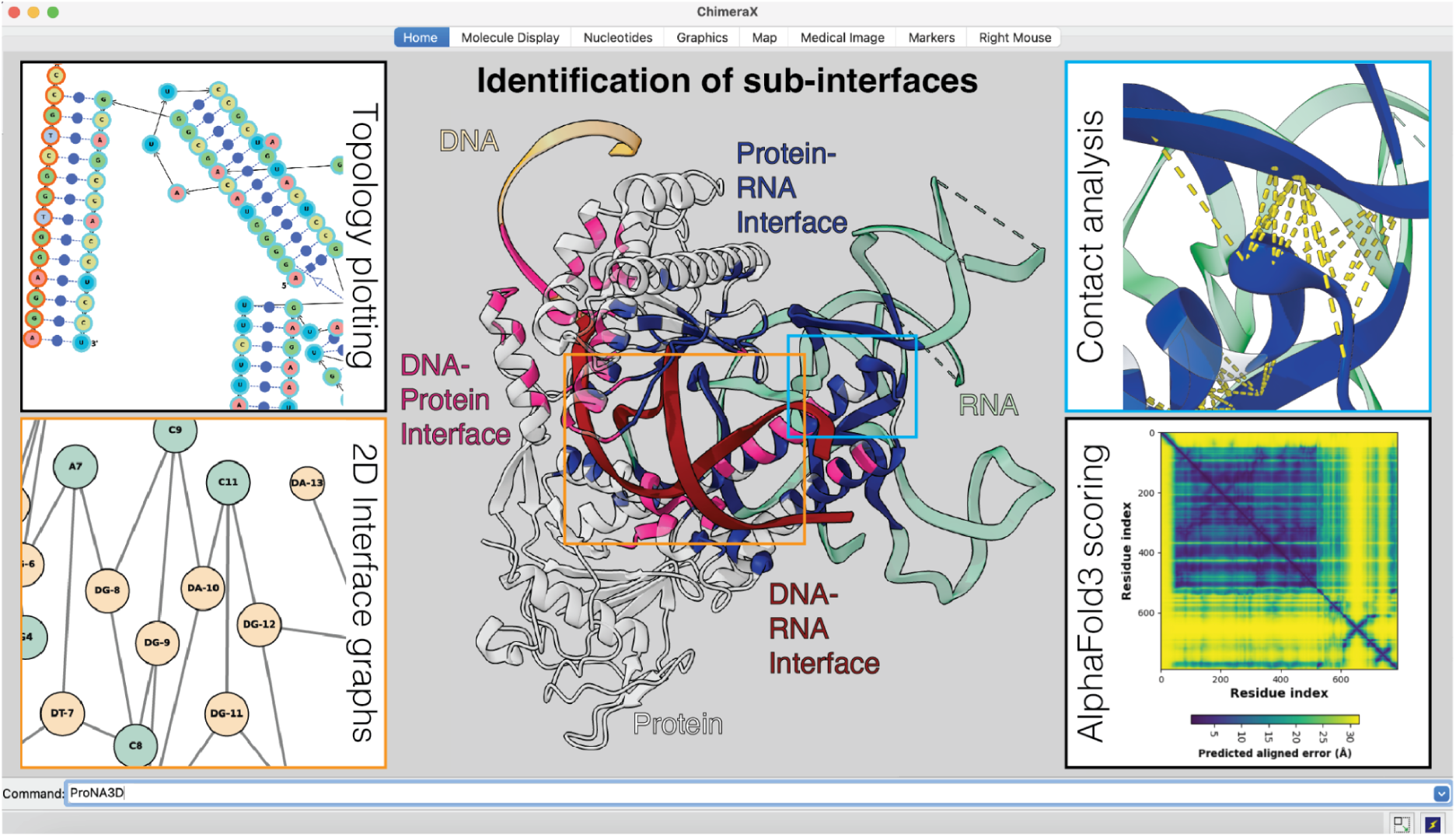
Overview of ProNA3D for integrated interface analysis of protein–nucleic acid complexes in ChimeraX. The central panel shows a representative protein-DNA-RNA complex (PDB ID: 9B0L), with distinct sub-interfaces automatically detected and annotated. The left panels illustrate ProNA3D’s interactive two-dimensional visualizations: a secondary-structure topology plot (top left box) and an interface graph depicting residues and nucleotides as nodes connected by interaction edges (bottom left, orange box). The right panels demonstrate ProNA3D’s interface contacts visualization (top right, blue box) and integration of AlphaFold3 confidence measures, e.g., PAE (bottom right box).

Moreover, application of the command-line version of ProNA3D to all high-resolution nucleic acid-containing interfaces in the PDB enabled large-scale structural characterization. The resulting analysis revealed distinct interface connectivity trends between nucleic acid-only and protein-nucleic acid interfaces, and highlighted specialized interaction modes such as base flipping.

## 3. Results

Below, we describe the application of ProNA3D to different types of complexes, including both experimentally determined (X-ray and cryoEM) and AI-based predicted complexes: X-ray structure of a dimeric hammerhead ribozyme RNA complex (PDB ID: 2OEU), a predicted human Fab antibody-HIV-1 RNA complex (Target PDB ID: 9C75), a predicted complex formed by the Cre recombinase from *Punavirus P1* bound to its target DNA (Target PDB ID: 7RHY), and the cryoEM structure of the *B. taurus* ATP-dependent DNA/RNA helicase DHX36 in complex with human G-quadruplex RNA (PDB ID: 8VV2, EMD-43547).

### 3.1. X-ray structure of an RNA-only complex: hammerhead ribozyme

To demonstrate the functionality of ProNA3D on experimentally determined structures, the plug-in was applied to a dimeric hammerhead ribozyme RNA complex (PDB ID: 2OEU). Varying the heavy-atom distance cutoff (d_HA_) revealed its strong impact on interface detection (Fig. 2A). At a d_HA_ of 3.0 Å, ProNA3D identified five sub-interfaces in the dimeric assembly. Increasing the d_HA_ to 4.0 Å reduced this to two sub-interfaces, and at 5.0 Å, only a single interface was identified. These observations highlight how d_HA_ defines the granularity of interface division.

**Figure 2:**
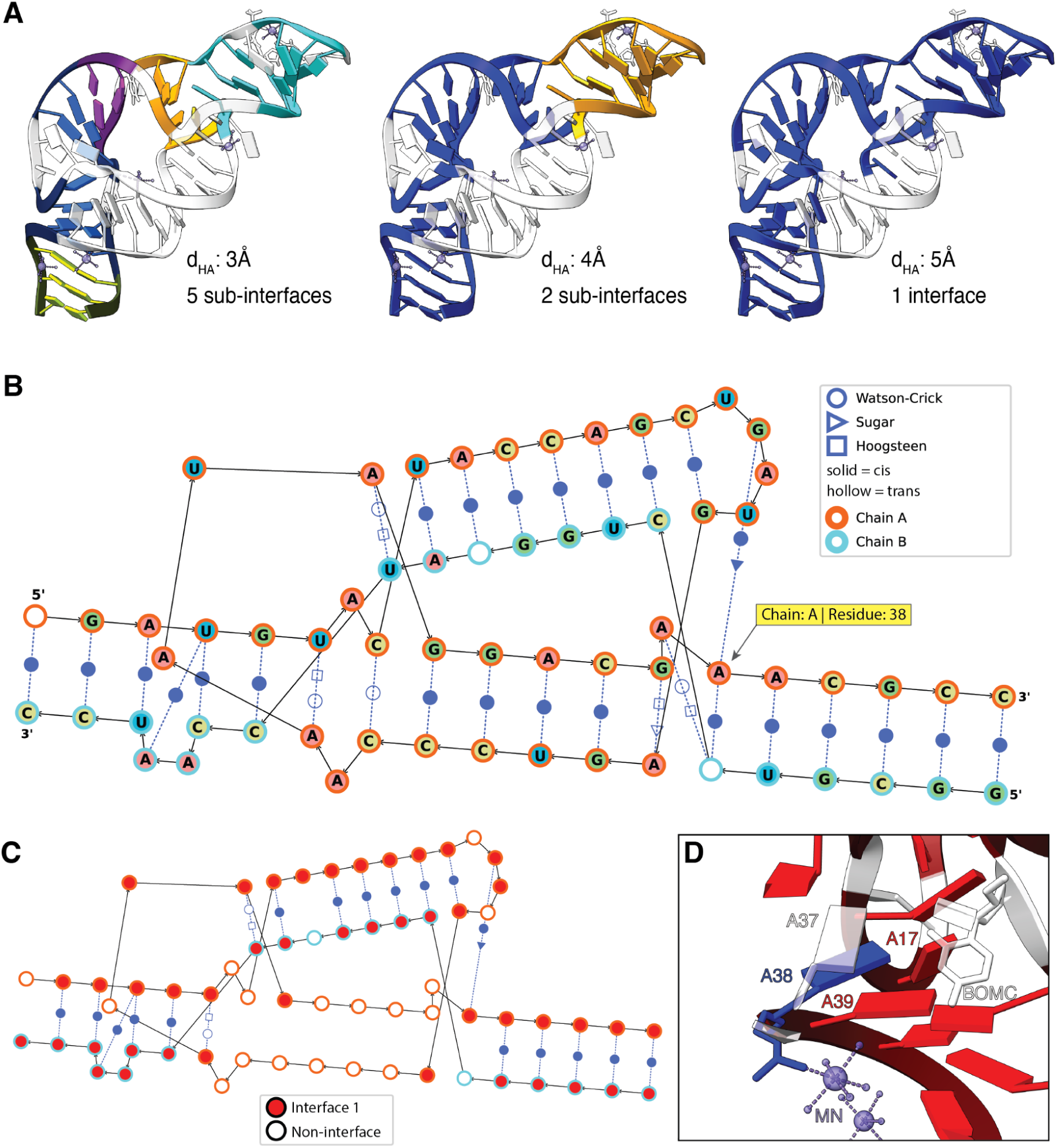
Influence of the heavy atom distance cutoff (d_HA_) and topology analysis of a dimeric hammerhead ribozyme RNA complex (PDB ID: 2OEU) using ProNA3D. **(A)** Sub-interfaces identified at d_HA_ of 3 Å (left), 4 Å (middle), and 5 Å (right). Nucleotides involved in inter-chain interfaces are shown in color, while non-interacting nucleotides are displayed in grey. **(B)** Topology plot of the ribozyme complex generated by ProNA3D using default coloring (A: red, U: blue, C: yellow, G: green). Black edges indicate backbone connectivity, and blue edges characterize base-pairing interactions annotated using the Leontis-Westhof classification. **(C)** Topology plot of the complex colored according to the detected inter-chain interface. **(D)** Close-up view of nucleotide A38 (blue) of chain A in the three-dimensional structure, highlighting its coordination of the Mn2+ ion.

Furthermore, the topology of the RNA-RNA complex was examined using Leontis-Westhof classification (Fig. 2B, C). ProNA3D detected 29 base-pair interactions, comprising nine intra-chain and 20 inter-chain pairs. The topology plot highlights nucleotide A38 in chain A as a central node in the interaction network (Fig. 2B). A38 forms backbone contacts with nucleotides 37 and 39, and pairs with nucleotide 17 through a cis Hoogsteen/Sugar-edge interaction. Moreover, it engages in a trans Watson-Crick/Hoogsteen base-pair with the 2′-O-methylated nucleotide at position 6 of chain B. This inter-chain interaction is shown to be part of the detected interface region (Fig. 2C). Inspection of A38 in the three-dimensional complex structure further shows that A38 coordinates a Mn^2+^ ion (Fig. 2D), illustrating that combining topology plots with structural visualization can highlight residues of potential structural importance for further investigation.

### 3.2. Predicted protein-RNA complex: human Fab antibody-HIV-1 RNA

To assess the performance of ProNA3D on structurally predicted complexes, we analysed the AlphaFold3 model of CASP16 target M1296, obtained from the CASP server (https://predictioncenter.org). The corresponding experimentally determined structure (PDB ID: 9C75) is formed by a trimeric assembly of the Fab heavy chain (H), light chain (L), and HIV-1 RNA (R, Rev Response Element stem-loop II G34U mutant) (Fig. 3A).

**Figure 3:**
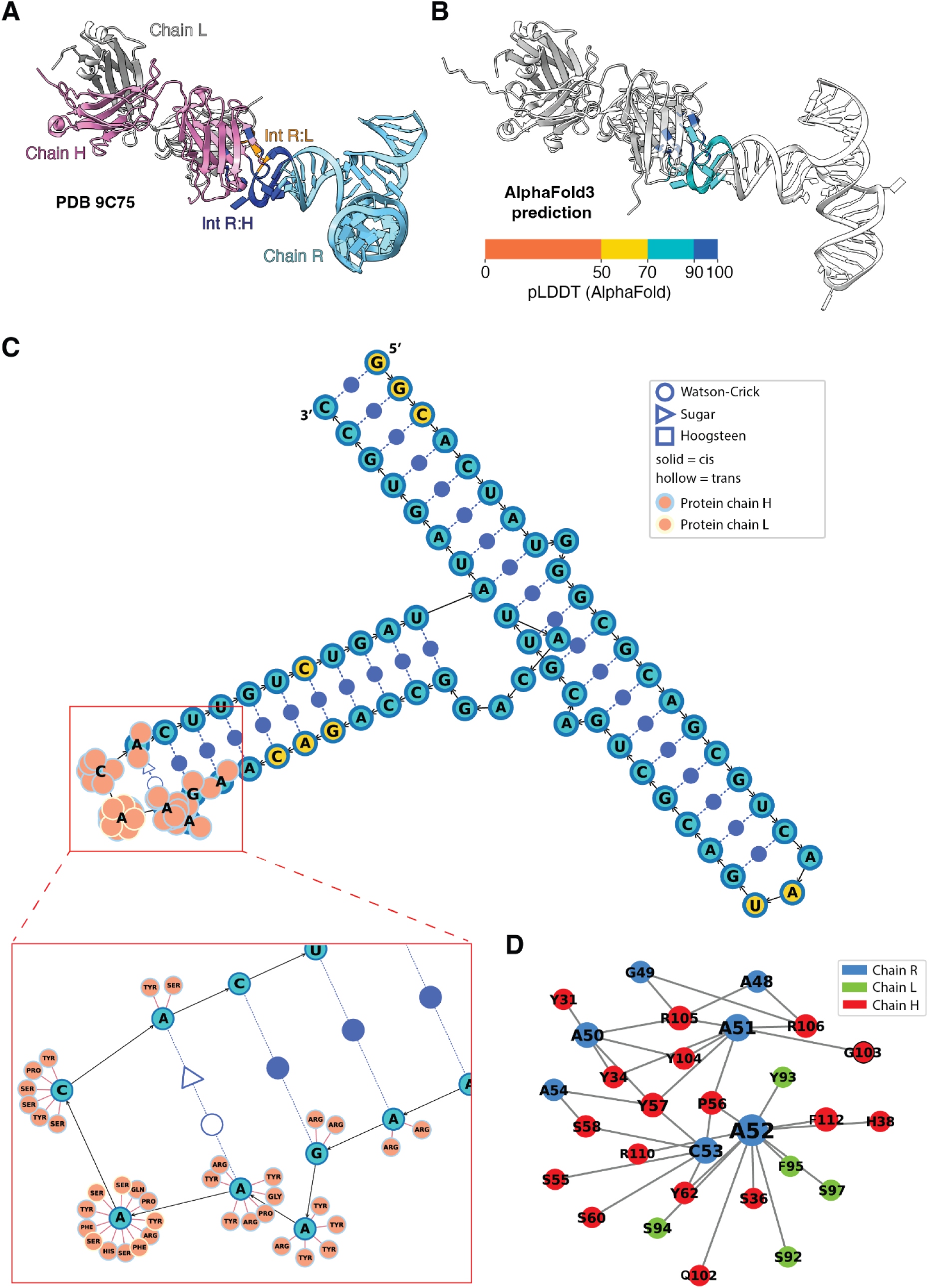
Analysis of the AlphaFold3 prediction for the CASP16 target M1296 in comparison with the experimentally determined structure (PDB ID: 9C75). **(A)** X-ray structure of the antibody–RNA complex with inter-chain interfaces identified by ProNA3D. The RNA (R) is shown in cyan, the Fab light chain (L) in grey, and the Fab heavy chain (H) in pink. Interface regions are highlighted in blue for the R-H interface and orange for the R-L interface. **(B)** AlphaFold3 prediction of the complex with interfaces colored by pLDDT. **(C)** Topology plot of the complex colored according to pLDDT values using ProNA3D. Black edges represent backbone connectivity, whereas blue edges indicate base-pairing interactions annotated using the Leontis-Westhof classification. Inset: The protein-interface regions of the RNA and interacting protein residues (salmon). **(D)** Two-dimensional interface network representation of the complex. Protein residues are in red (Chain A) and green (Chain B), and RNA is in blue (Chain C).

The analysis of the complex with ProNA3D identified interfaces between the RNA and both heavy and light antibody chains in the predicted model (Fig. 3B). The R-H interface involved 23 interacting residues and nucleotides, whereas the R-L interface comprised five interaction partners. Structural comparison of the AlphaFold3 prediction and the experimental structure showed highly similar orientations of the two protein chains, while the global fold of the RNA differed slightly (Fig. 3A, B). However, the interface pattern detected by ProNA3D was largely conserved between the two structures, as reflected by an Interface Contact Similarity (ICS) score of 0.875 [31,32].

Analysis of the predicted local distance difference test (pLDDT) [33] scores of the predicted interfaces indicated high confidence in the interacting regions, with most residues exhibiting pLDDT values between 70 and 90 and several exceeding 90 (Fig. 3B). Residue-level comparison of interfaces identified in the predicted and experimental structure revealed that only one residue in the R-L interface (L97) and one nucleotide in the R-H chain interface (R47) differed between the two models.

To further analyse the predicted complex, ProNA3D was used to generate a topology plot of the AlphaFold3 model, incorporating pLDDT-based node coloring and visualization of protein-RNA contacts (Fig. 3C). The secondary structure representation showed that both antibody chains interact with the tip region of the RNA stem, which contains a bulge within the interface while being characterized by consistent base-pairing.

ProNA3D also provides a two-dimensional interface representation in which interacting residues and nucleotides are displayed as nodes linked by edges. Examination of this network for the RNA-antibody complex highlighted nucleotide A52 as a highly connected node, suggesting it as a candidate for further structural and functional investigation (Fig. 3D).

### 3.3. Predicted protein-DNA complex: Cre recombinase from *Punavirus P1* bound to its target DNA

Next, we analyzed a predicted protein-DNA complex and evaluated the impact of different pLDDT thresholds on interface definition. To this end, we applied ProNA3D to the AlphaFold3 prediction of a dimeric complex formed by a triple-point mutant of Cre recombinase from *Punavirus P1* bound to its target DNA (Target PDB ID: 7RHY; Suppl. Fig. 1A). Using the default d_HA_ and distance for the interface clustering (d_C_) and applying a pLDDT cutoff of 50, ProNA3D identified two sub-interfaces comprising 102 and 5 interacting residues and nucleotides, respectively (Suppl. Fig. 1B). For this threshold, the calculated interface pLDDT (ipLDDT) [30] values were 88.51 for the sub-interface 1 and 89.62 for the sub-interface 2, while the corresponding interface pairwise alignment error (iPAE) [30,34] values were 2.66 Å and 2.26 Å, respectively.

Raising the pLDDT cutoff to 70 reduced the size of sub-interface 1 to 95 interacting residues and nucleotides, whereas sub-interface 2 remained unchanged (Suppl. Fig. 1C). Under these conditions, ipLDDT values of 89.43 and 89.62 were calculated for sub-interfaces 1 and 2, respectively, as well as iPAE values of 2.41 Å and 2.26 Å. Further increasing the cutoff to 90 resulted in substantially smaller interfaces comprising 16 (sub-interface 1) and 3 (sub-interface 2) interacting residues and nucleotides (Suppl. Fig. 1D). For this threshold, ipLDDT values increased to 90.37 and 90.29, and iPAE values decreased to 2.08 Å and 2.05 Å for the two sub-interfaces. Notably, interfaces defined using a pLDDT cutoff of 90 correspond to highly reliable regions, which makes them particularly suitable for downstream applications.

Other confidence metrics remained constant across all interface definitions, as these values are not recalculated by ProNA3D. These included a pTM score of 0.92, an ipTM score of 0.89, and a *model confidence* of 0.90, which further supported the overall high quality of the predicted complex.

### 3.4. cryoEM structure of a protein-RNA complex

*B. taurus* ATP-dependent DNA/RNA helicase DHX36 / human G-quadruplex RNA Large RNA-or DNA-containing macromolecular assemblies, such as ribozymes, are increasingly solved using cryoEM. To facilitate interface-focused analysis of cryo-EM-derived complexes, ProNA3D includes a feature that allows users to directly zone into the density corresponding to identified interface regions.

We used this feature to analyse the ATP-dependent DNA/RNA helicase DHX36 from *B. taurus* bound to a human G-quadruplex RNA (cMyc G-quadruplex, class I; PDB ID: 8VV2) at a reported resolution of 2.6 Å (Fig. 4A). Applying default parameters for d_HA_ and d_C_, ProNA3D detected two sub-interfaces (Fig. 4B, C). Zoning the density around sub-interface 1 revealed a high local resolution, indicating reliable modeling of both backbone and side-chain interactions (Fig. 4B, D). By contrast, sub-interface 2 exhibits lower local resolution (Fig. 4C, D), suggesting that modeled side chains in this region should be interpreted cautiously. Notably, residues Lys854 and Lys855 in sub-interface 2 are only partially modeled in the deposited structure, likely reflecting the reduced local resolution in this area (Fig. 4D). However, ProNA3D identified these residues as interface participants, supporting their proposed placement and potential interaction with the RNA, consistent with the results from Banco et al. [35]. Analysis of the complex predicted by AlphaFold3 further revealed that the pLDDT profile of the prediction agrees with the local resolution pattern (Fig. 4E), with sub-interface 2 displaying lower predicted confidence than sub-interface 1. The reduced local resolution and low confidence of the model in the region may reflect increased structural flexibility. This example demonstrates that interface-based zoning with ProNA3D can provide an effective approach for examining protein-nucleic acid interactions in the context of cryo-EM maps, which often suffer from heterogeneous resolution.

**Figure 4:**
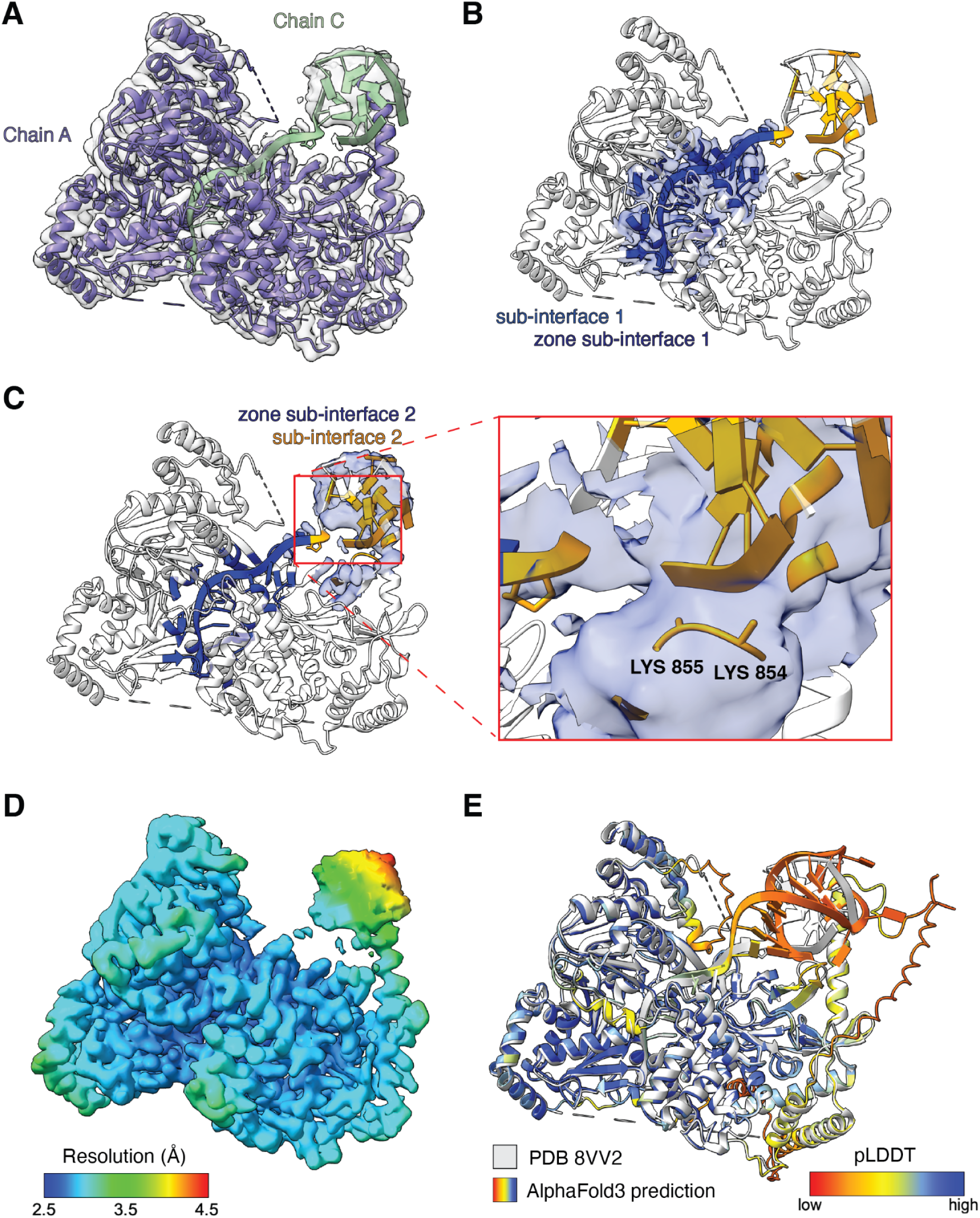
ProNA3D analysis of the Cryo-EM structure of the ATP-dependent DNA/RNA helicase DHX36 in complex with G-quadruplex RNA (EMD: PDB: 8VV2, EMD-43547). **(A)** An overall view of the dimeric complex, with the helicase shown in purple, RNA in green, and the density map in grey. **(B)** Sub-interfaces identified by ProNA3D, with sub-interface 1 in blue and sub-interface 2 in orange. The density map (blue) is zoned around sub-interface 1. **(C)** The density map (blue) is zoned around sub-interface 2. Inset: Lys854 and Lys855, which are not fully modelled (due to lack of density in this region) but participate in the interface (sub-interface 2), are highlighted. **(D)** Local resolution (Å) map of the complex, with blue indicating better resolution and red indicating worse resolution. The map was calculated using RELION-4.0.0 [36], with a mask applied to the modeled region. **(E)** Superposition of the AlphaFold3 prediction of the complex, colored by pLDDT (red indicating low values and blue indicating high values, helicase residues: 1-972, RNA nucleotides: 1-27) on the cryoEM corresponding deposited structure (PDB ID: 8VV2) (grey, helicase residues: 15-948 with unresolved regions not modeled, RNA nucleotides: 2-27 with unresolved regions not modeled).

### 3.5. Systematic analysis of protein-nucleic acid and nucleic acid-only interfaces using ProNA3D

In addition to a graphical implementation, ProNA3D is available as a command-line tool, enabling rapid, large-scale analysis of macromolecular complexes without visualization. Here, the command-line version of ProNA3D (default value of 5 Å for both d_HA_ and d_C_) was applied to all nucleic acid-containing complexes in the PDB (for the protocol for dataset curation, see Methods). These were classified into RNA-RNA (n = 4,797), RNA-DNA (n = 450), DNA-DNA (n = 4,590), protein-DNA (n = 11,791), and protein-RNA (n = 23,616) interfaces.

To characterize structural differences between interface classes, we analysed residue and nucleotide connectivity. Connectivity is defined per residue or nucleotide as the number of distinct residues or nucleotides from the interaction partner that contact that given residue/nucleotide across the interface. For each interface, the maximum observed connectivity (whether nucleotide or residue) was recorded, and connectivity distributions were examined both at the interface level and at the level of individual residue and nucleotide types. Moreover, residue-and nucleotide-type occurrence distributions at the interface were analysed.

#### 3.5.1. Maximum connectivity distributions for different interface classes

Nucleic acid-only interfaces exhibited highly consistent maximum connectivity values across interaction types. For each interface, connectivity was calculated for every residue or nucleotide at the interface, and the highest observed value was defined as the maximum connectivity of that interface. RNA-RNA, RNA-DNA, and DNA-DNA interfaces all showed median maximum connectivity values of 3 interaction partners per nucleotide, with relatively low variability (RNA-DNA: σ = 0.76; DNA-DNA: σ = 0.88) (Fig. 5, Suppl. Table 2). RNA-RNA interfaces displayed slightly broader distributions (σ = 1.83), consistent with the greater structural diversity and tertiary interactions characteristic of RNA assemblies. The local maximum in the DNA-DNA interaction distribution at a connectivity of 1 could be explained in the majority of cases by nicked DNA structures [37] (e.g., from endonuclease cleavage), as exemplified for the DNA substrate for DNA repair enzymes (PDB-ID: 1VTE, Fig. 6A). Although the phosphodiester bond is broken, base stacking across the nick can be retained, producing non-covalent interactions at connectivity of 1 (Fig. 5, DNA-DNA). In contrast, protein-nucleic acid interfaces showed higher maximum connectivity values and substantially broader distributions (protein-RNA: σ = 3.32; protein-DNA: σ = 3.48) (Fig. 5, Suppl. Table 2). Protein-DNA and protein-RNA interfaces exhibited median maximum connectivity values of 5 and 4, respectively, corresponding to an increase of approximately 2 and 1 additional interaction partners compared to nucleic acid-only interfaces. The large standard deviations indicated a heterogeneous interface organization, reflecting the diversity of protein-nucleic acid binding modes and the smaller size of amino acid residues in comparison to nucleic acids [38].

**Figure 5:**
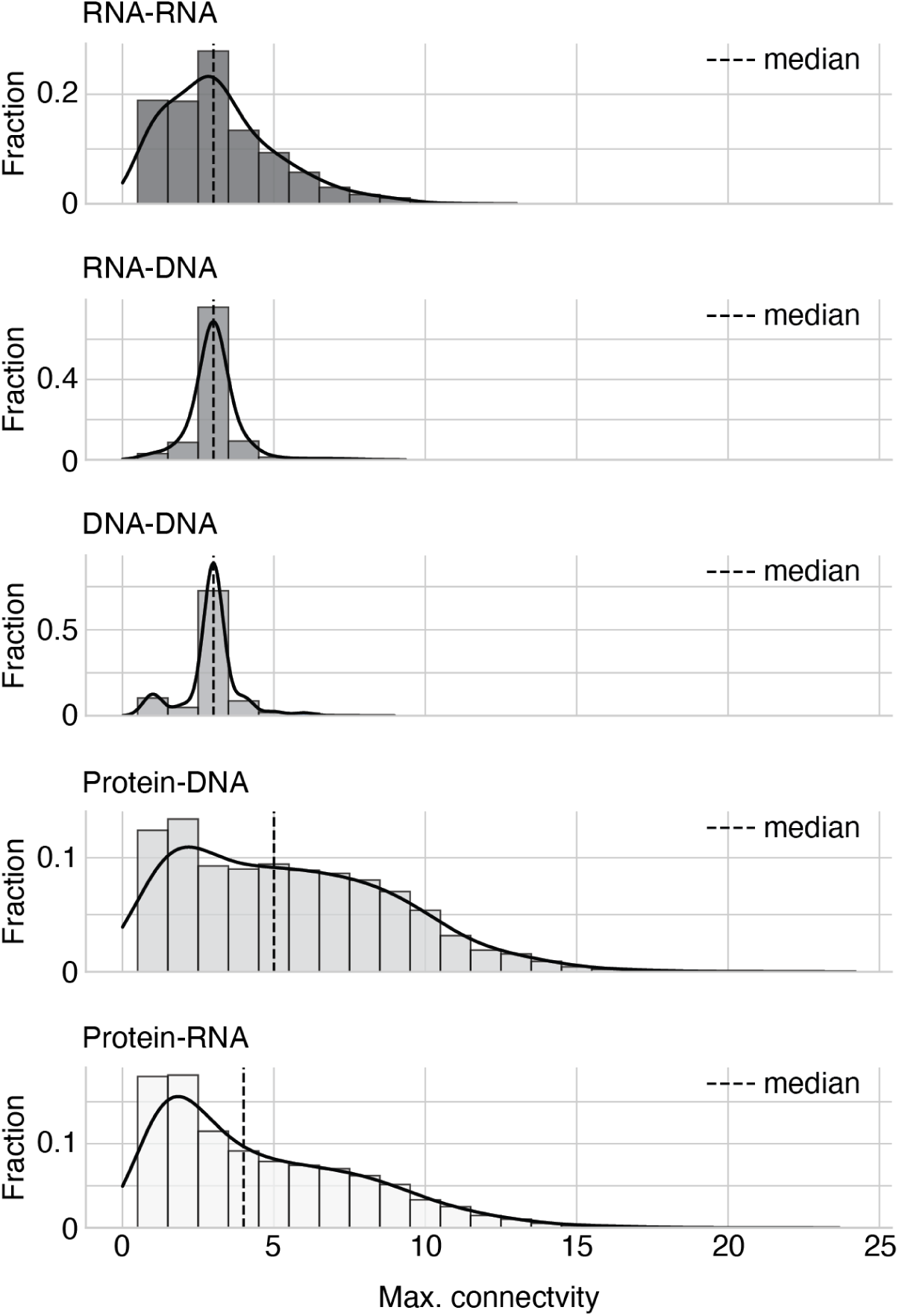
Maximum connectivity across nucleic acid-containing interface types. Connectivity was defined as the number of distinct residues or nucleotides contacted by a residue or nucleotide across an interface. For each interface, the highest connectivity value was extracted. Shown are RNA-RNA, RNA-DNA, DNA-DNA, protein-DNA, and protein-RNA interfaces. Median values are indicated for each interface class (dotted line).

**Figure 6:**
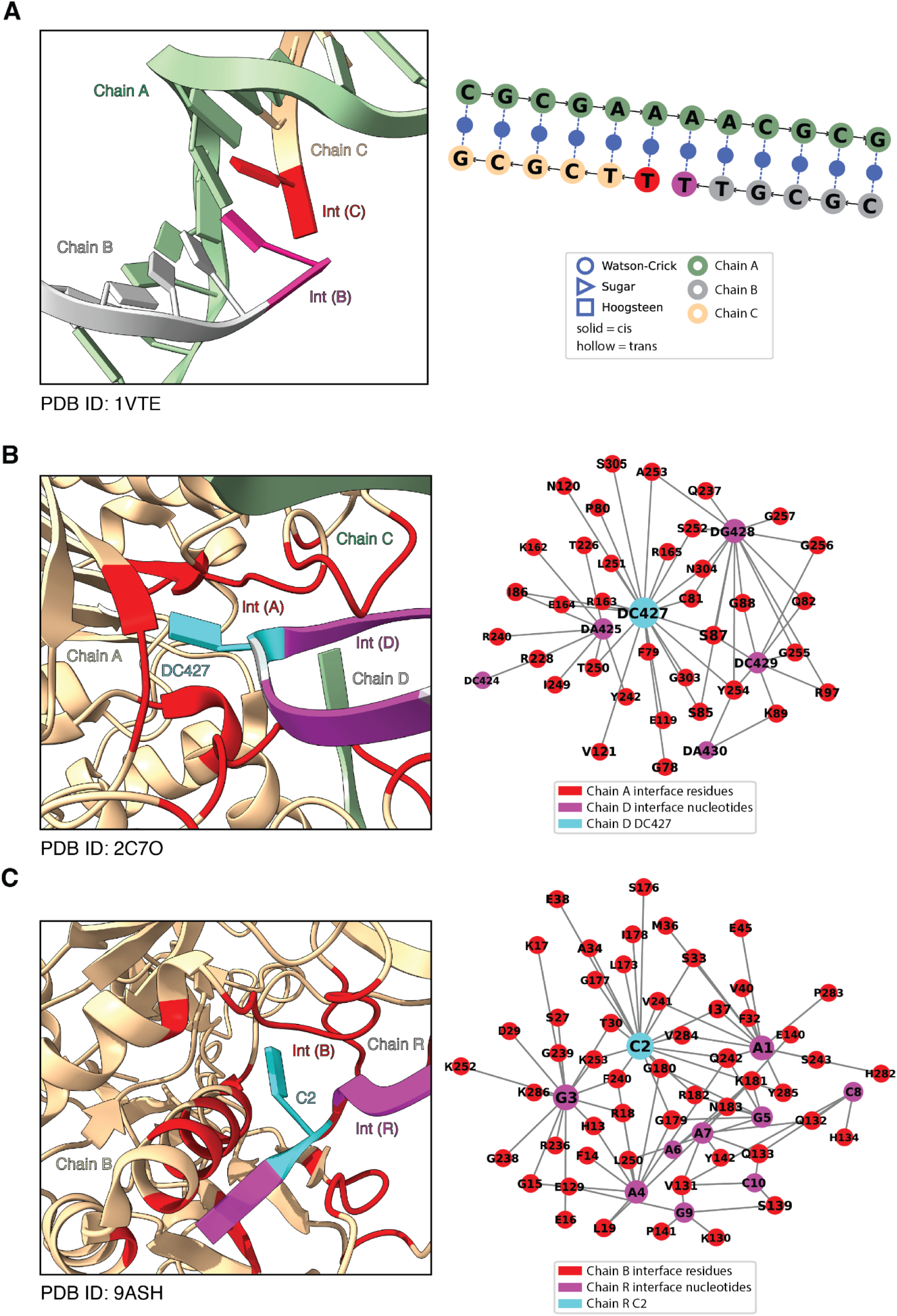
Examples illustrating nicked DNA (low-connectivity outliers) and nucleotide base flipping associated with high connectivity (≥18). Interfaces of the highlighted complexes are shown, with interface residues from interaction partner 1 in red and from interaction partner 2 in magenta, except the flipped nucleotides, which are shown in cyan. Corresponding secondary structure plot (A) and interaction networks (B,C) derived from ProNA3D are displayed for each complex. **(A)** Nicked DNA duplex (PDB ID: 1VTE), illustrating base stacking across a nicked DNA strand. **(B)** DNA methyltransferase (Chain A, beige) from *H. haemolyticus* bound to its target DNA (Chain D, grey) (PDB ID: 2C7O). **(C)** CRISPR-associated protein Csm4 (Chain B, beige) from *L. lactis* in complex with crRNA (Chain R, grey) (PDB ID: 9ASH).

Pairwise comparisons of maximum connectivity distributions confirmed that the differences between nucleic acid-only and protein-nucleic acid interfaces are statistically significant across interface classes (Mann-Whitney U test, Suppl. Table 3). Not surprisingly, the differences between different types of nucleic acid-only interfaces are not significant.

Moreover, repeating the analysis using threshold pairs of 4/5 Å, 4/4 Å, and 5/4 Å to variations in d_HA_ and d_C_, resulted in similar trends for the maximum-connectivity distributions (Suppl. Fig. 2; Suppl. Tables 4-6). Although stricter cutoffs reduced absolute connectivity values, protein-DNA and protein-RNA interfaces consistently exhibited higher mean maximum connectivity and broader distributions than nucleic acid-only interfaces. The differences in median maximum connectivity were less pronounced, however, the overall patterns in distributions remained unchanged, indicating that the trends observed for the 5 Å threshold for both parameters are robust to the choice of distance cutoff.

Protein-nucleic acid complexes contained a subset of cases with exceptionally high connectivity. To investigate these outliers structurally, interfaces with connectivity ≥ 18 were examined in detail. In most cases, these involved enzymes interacting with DNA, such as methyltransferases (typically connectivity ≥ 20), where individual bases were flipped out of the helix for covalent modification or excision (e.g., PDB: 2C7O) [39]. In these structures, the flipped nucleotide was almost completely enclosed by surrounding protein residues, as illustrated for the DNA methyltransferase from *H. haemolyticus* bound to its target DNA, resulting in exceptionally high single-nucleotide connectivity (Fig. 6B). Many interfaces with connectivity values between 18 and 20 involved components of the CRISPR-Cas machinery, as exemplified for the CRISPR-associated protein Csm4 from *L. lactis* that interacts with the crRNA (Fig. 6C, PDB-ID: 9ASH). In this structure, C2 of the RNA is flipped out with a connectivity of 18. This binding mode appeared to be conserved, as we also observed a highly-similar flipped conformation and high connectivity for Csm4 from *S. thermophilus* (PDB-ID: 9NO4, Suppl. Fig. 3). This observation is consistent with previous structural analyses, which describe nucleotide flipping at the 5′ tag induced by a β-hairpin of Csm4 [40].

#### 3.5.2. Individual residue-and nucleotide-level connectivity for different interface classes

Individual residue-and nucleotide-level analysis further highlighted the differences between nucleic acid-only and protein-nucleic acid complexes (Suppl. Fig. 4,5, Suppl. Table 7-14). DNA-DNA interfaces showed tightly constrained connectivity patterns across all four nucleotides with small standard deviations (Suppl. Fig. 4, Suppl. Table 7), whereas RNA-RNA interfaces exhibited broader distributions, with adenine and guanine showing slightly elevated connectivity (Suppl. Fig. 4, Suppl. Table 8). RNA-DNA interfaces displayed similar connectivity behavior for RNA and DNA nucleotides, with only adenine and thymine showing higher median connectivity of 3, consistent with geometry-constrained hybrid base pairing (Suppl. Fig. 4, Suppl. Tables 9,10).

In protein-RNA and protein-DNA interfaces, connectivity varied strongly by amino acid type, with positively charged residues, particularly arginine, showing the highest connectivity values, identifying them as consistent interaction hot spots (Suppl. Fig. 5, Suppl. Tables 11-14).

#### 3.5.3. Occurrence distributions of residue-and nucleotide-types for different interface classes

Occurrence analyses of residues and nucleotides mirrored the already described trends. DNA-DNA interfaces exhibited uniform nucleotide frequencies (Suppl. Fig. 6, Suppl. Table 7), while in RNA-RNA and RNA-DNA interfaces purines were slightly more abundant than pyrimidines (Suppl. Fig. 6, Suppl. Table 8-10). Protein-RNA interfaces were enriched in guanine and adenine, and protein-DNA interfaces in adenine and thymine (Suppl. Fig. 6, Suppl. Tables 11,13). On the protein side, the positively charged arginine and lysine were most frequent, followed by polar residues, with hydrophobic residues underrepresented, aligning with the observations from [41] (Suppl. Fig. 7, Suppl. Tables 12,14).

Overall, this large-scale analysis demonstrated that nucleic acid-only interfaces are more defined by constrained and uniform interaction geometries, whereas protein-nucleic acid interfaces exhibit greater heterogeneity.

## 4. Discussion

In this work, we present ProNA3D, a new tool that enables the identification and analysis of interfaces in protein-nucleic acid, and nucleic acid-only complexes. ProNA3D identifies inter-chain interfaces based on spatial properties and further subdivides them into distinct sub-interfaces, facilitating a more detailed characterization of complexes.

ProNA3D extends interface analysis beyond three-dimensional contacts by providing analysis options for nucleic acid and protein-nucleic acid complexes, including topology plots, two-dimensional interface graphs, and predicted interface-specific confidence metrics. Together with support for analysing the interfaces in the context of cryo-EM density maps, these features enable assessment of interaction quality and structural reliability in both predicted and experimental complexes.

Several computational tools are available for analyzing protein-nucleic acid interactions, including DNAproDB [19], RNAproDB [20], and RNAView [27], as well as interface analysis platforms such as PLIP [25] and VoroContacts [26]. However, these tools are primarily tailored to experimentally resolved structures, often focus on either proteins or nucleic acids, and lack unified, interactive workflows for mixed complexes. ProNA3D addresses this gap by allowing for the analysis of both experimental and predicted nucleic acid-containing complexes. Furthermore, in addition to being available as a command line tool, it is also available as a ChimeraX plug-in, providing an integrated, graphical environment for the analysis of such assemblies.

By enabling detailed interface inspection, ProNA3D facilitates the identification of highly connected interface residues and nucleotides, providing a framework for exploring interaction patterns and prioritizing candidates for subsequent experimental validation, including mutational analyses. This capability is illustrated by our analyses of a dimeric hammerhead ribozyme RNA complex (PDB ID: 2OEU) and a predicted human Fab-antibody-HIV-1 RNA complex (target PDB ID: 9C75), in which nucleotides A38 and A52, respectively, were identified as highly connected interaction nodes and inferred to play structurally important roles.

Recent advances in AI-driven structure prediction have substantially expanded the structural coverage of nucleic acid-containing complexes by allowing access to assemblies difficult or impossible to characterize experimentally. As a result, predicted complex models are becoming increasingly important resources, highlighting the need for tools that support their detailed and reliable analysis. While existing platforms such as RNAproDB and DNAproDB allow the analysis of user-provided structures, they are primarily tailored to experimentally resolved models and do not support the visualization of prediction confidence metrics in a structural context, limiting their applicability to predicted complexes. ProNA3D addresses this limitation by integrating interface-specific confidence measures, including ipLDDT and iPAE, directly into the structural analysis workflow, allowing users to assess prediction quality directly at the interaction level. This capability facilitated, for example, the identification of high-confidence sub-interfaces in a predicted complex between the *Punavirus P1* Cre recombinase and its target DNA (PDB ID: 7RHY), illustrating the utility of ProNA3D for evaluating predicted complexes.

Furthermore, in recent years, cryoEM has become the primary method of choice for determining the structures of nucleic acid-containing complexes. However, cryoEM maps often show strong local variations in resolution, which can complicate structural interpretation. Atomic models derived from these maps do not always capture local variations that are often associated with molecular flexibility or functional dynamics at interface regions [42]. By allowing density maps to be zoned around detected interface regions and combined with model-derived confidence scores, such as b-factors or pLDDT, ProNA3D supports local assessment of model quality in large assemblies. We demonstrated this approach on the cryo-EM structure of the *B. taurus* ATP-dependent DNA/RNA helicase DHX36 in complex with human G-quadruplex RNA (PDB ID: 8VV2; EMD-43547), allowing for focused analysis of specific interface regions suffering from lower local resolution.

The large-scale connectivity analysis enabled by ProNA3D not only confirmed established principles of nucleic acid interfaces but also revealed subtle and functionally relevant variations. The relatively uniform connectivity of nucleic acid-only interfaces reflects the regularity of duplex DNA and the structured tertiary contacts in RNA [43,44]. In contrast, the broader and more heterogeneous connectivity patterns in protein-nucleic acid interfaces highlight the diversity of protein-nucleic acid interaction modes [38].

Rare high-connectivity outliers, including base-flipping in DNA methyltransferases and CRISPR-Cas Csm4 proteins, emphasize specialized interactions beyond typical interface metrics. The conservation of flipped nucleotides across species suggests that such extreme connectivity may be linked to structural and functional importance. Residue-and nucleotide-level trends, such as the enrichment of positively charged residues at protein interfaces, align with known electrostatic and geometric constraints [41], whereas features like the DNA-DNA maximal connectivity of 1, indicating nicked DNA, illustrate the ability of ProNA3D to capture also subtle structural variations.

Overall, ProNA3D provides an interactive framework for the comprehensive analysis of both experimentally determined and computationally predicted structures of protein and nucleic acid complexes and supports a wide range of downstream structural and functional studies. Integration within ChimeraX further extends analytical possibilities through access to other analysis tools from the ChimeraX suite.

In the future, the combination of 3D interface identification, topology plots, and interface graphs in ProNA3D allows the framework to support both detailed case-specific studies and large-scale structural analyses, enabling the systematic discovery of recurring interaction and secondary-structure patterns, connectivity signatures, and interaction features. However, high connectivity should be regarded as a hypothesis for further investigation rather than direct indicators of functional importance. An important limitation of the current ProNA3D implementation is the lack of automatic support for modified nucleotides and modified amino acids. As chemically modified nucleotides are prevalent in many experimentally determined RNA structures and can influence RNA secondary structure formation [45,46], incorporating automated handling of these modifications will be part of future development of the software. Furthermore, while current secondary structure annotation in ProNA3D relies on DSSR, future implementations could expand compatibility with additional secondary structure annotation approaches, such as RNAView [27], to increase flexibility and applicability across diverse RNA structures and analysis workflows.

## 5. Methods

### 5.1. Programming languages and general tools

ProNA3D is developed in Python 3.9 and uses a front-end built with HTML, JavaScript, Qt, and Jinja2 to enable an interactive user interface within the application. The tool is distributed as a plug-in for UCSF ChimeraX version 1.10 [29], while still maintaining compatibility with previous releases of the software.

### 5.2. General parameters and functions

ProNA3D relies on the interface detection strategy originally introduced in PICKLUSTER [30], but substantially broadens its scope and functionality. While PICKLUSTER was designed specifically for protein-protein interfaces, ProNA3D generalizes the approach for the analysis of interaction sites across protein-RNA, protein-DNA, RNA-RNA, RNA-DNA, and DNA-DNA complexes.

It is worth noting that ProNA3D is not able to handle modified nucleotides and protein residues. In order to include such entities in the interface calculation, it is possible to modify them manually in ChimeraX.

The plug-in operates directly on PDB or mmCIF structure files loaded into ChimeraX, and can be applied to both experimentally determined structures and predicted models (e.g, AlphaFold3). Once the calculation is started, ProNA3D automatically identifies “sub-interfaces” of contacting residues or nucleotides and reports the results in the ChimeraX Log.

To accommodate different analysis objectives, the plug-in provides three modes for the detection of interfaces:

1. **Pairwise interface detection** between user-selected chains, designed for situations where the interaction between two specific components is of interest.
2. Interface detection across **the entire complex**, offering a global view of all inter-chain contacts within multimeric complexes of high stoichiometry.
3. Interface detection for a **single chain against all other chains**, which is particularly helpful when the focus is on the interaction landscape of one specific component.

ProNA3D can additionally report specific interaction partners on the residue/nucleotide-level, which is valuable for identifying positions involved in multiple interfaces (“interaction hubs”). For downstream analyses, the tool allows exporting all detected interfaces to a CSV file, enabling further processing or visualization with external software.

### 5.3. Interface extraction

Interacting residues are identified by calculating distances between their heavy atoms, using a KDTree data structure from *SciPy* [47]. The heavy-atom distance cutoff (d_HA_), which defines interacting atoms, can be selected by the user in ProNA3D. However, ProNA3D applies a default d_HA_ of 5.0 Å [34].

The interface *I* is calculated as follows:

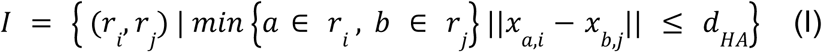

where *r_i_* and *r_j_* are residues (or nucleotides) on chains *i* and *j*, *a* and *b* denote their corresponding heavy atoms with coordinates *x*, and *d_HA_*is the heavy-atom distance cutoff.

Users can further decide whether to apply the interface clustering algorithm provided by ProNA3D. When enabled, the plug-in groups interacting residues or nucleotides into “sub-interfaces” using a centroid-based approach. For each residue or nucleotide participating in the inter-chain interface, the centroid of all atoms is computed. If the distance between these centroids is below a user-specified cutoff distance *d_C_* (default is 5.0 Å), they are clustered into the same sub-interface *S_k_*, where the set of sub-interfaces *S* is defined as:

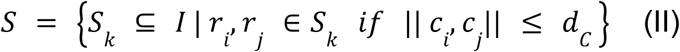

where *r_i_*, *r_j_* are residues or nucleotides within the interface *I*. The centroids *c_i_* and *c_j_* of a residue, or a nucleotide, *r_i_* and *r_j_*, on chains *i* and *j*, are defined as the mean position of their atoms, and two elements belong to the same sub-interface if the distance between their centroids is less than, or equal to, the cutoff *d_C_*.

ProNA3D supports an additional feature, aimed at the analysis of interfaces of structural models, where some regions may suffer from low confidence. To exclude such regions, users can specify a confidence score threshold that is based on the values per residue (averaged from the corresponding per-atom values) from the B-factor column (e.g., pLDDT [33]). Residues and nucleotides displaying a pLDDT below the selected cutoff are then excluded from the computation of the interface.

### 5.4. Topology and 2D interface plotting

In ProNA3D, the secondary structure of nucleic acids is displayed as topology plots, which represent the overall fold of an RNA or DNA molecule in a simplified two-dimensional graph [27].

To generate topology plots, ProNA3D uses the program DSSR (Defining the Secondary Structure of RNA) [21]. The analyses presented in this manuscript were performed using DSSR v2.6.0. DSSR detects all base pairs in a structure, identifies canonical and non-canonical backbone features, and captures complex motifs such as pseudoknots. When a topology plot is requested, ProNA3D executes DSSR, stores the resulting output to a dedicated directory, and retrieves the generated JSON file, which contains the secondary-structure information required for visualization. The subsequent parsing and transformation of the DSSR JSON file follow an approach adapted from the source code of the RNAscape webserver [22]. Visualization of the final plot is performed using the Matplotlib [48] and NetworkX [49] packages.

For the base-pairing annotation used in the topology plot, ProNA3D offers four options: (1) the Leontis-Westhof classification (12 classes) [50]; (2) the Saenger classification (28 classes) [51]; (3) DSSR’s own annotation scheme (closely related to Leontis–Westhof) [21]; and (4) a mode without base-pairing annotations.

Interacting residues and nucleotides are depicted as nodes linked by edges, and an interactive selection that highlights the corresponding elements in the 3D structure is supported. Black edges indicate backbone connectivity, and blue edges characterize base-pairing interactions.

To enable the link between the detected three-dimensional interface and the secondary-structure visualization, ProNA3D provides the option of coloring the topology plot according to the sub-interfaces. For predicted models with a confidence score in the B-factor column (such as AlphaFold3 pLDDT), the topology plot can also be colored by these values, providing a visual indication of model confidence in the context of secondary structure.

For protein-nucleic acid complexes, protein-nucleic acid contacts can be added to the topology plot, with interacting residues being linked to their corresponding nucleotide nodes. To preserve readability, each protein residue is linked to only a single nucleotide node, which may result in the duplication of protein residue nodes when necessary.

The topology plot is fully interactive, enabling manual adjustment of node positions and selection of residues or nucleotides in the three-dimensional structure directly via the plot.

Since the topology plotting relies on DSSR, which requires a user license, the path to the DSSR executable (x3dna-dssr) must be provided. While ProNA3D was tested with DSSR v2.6.0, compatibility with other DSSR versions may depend on the output format generated by the respective release. Future DSSR versions will be tested for compatibility. For users without access to DSSR, or when only a simplified representation is required, ProNA3D offers a two-dimensional plot of the identified complex interface. In that 2D plot, node size is correlated with its level of connectivity. Visualization is performed utilizing the Matplotlib and NetworkX packages.

### 5.5. AlphaFold3 scoring

ProNA3D includes a set of scoring metrics designed for assessing protein complex models produced by AlphaFold3. Among these is the calculation of the ipLDDT (interface predicted local distance difference test), an interface-specific version of the pLDDT [33]. For this, ProNA3D averages the atom-level values across all positions that form the interface, offering an estimate of the local confidence in the predicted contacts. The ipLDDT is calculated as follows:

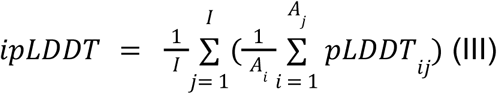

where *I* is the number of interface residues or nucleotides, *A* is the number of atoms in residue *j*, and *pLDDT_ij_* is the pLDDT of the atom *i* in residue *j*.

Moreover, the plug-in provides the option of computing the iPAE (interface predicted aligned error) [30,52]. To calculate the iPAE, all PAE [34] entries associated with the interface residues and nucleotides are extracted from the PAE matrix and then averaged to obtain a single score for each interface or sub-interface. The iPAE is calculated as follows:

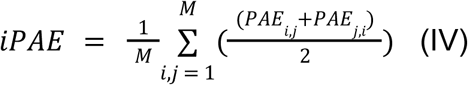

where *PAE_i_*_,*j*_ and *PAE_j_*_,*i*_ denote the two entries of the PAE matrix corresponding to residue pairs involved in the interface *I*, and *M* is the total number of residue pairs on chains *i* and *j* (each corresponds to two PAE entries).

In addition, the plug-in reports other AlphaFold3 confidence metrics, such as pTM [33], ipTM [34], *model confidence* (MC = 0.8*ipTM + 0.2*pTM), the maximum PAE, and the PAE matrix. Except for ipLDDT, these metrics require the scoring metric JSON files generated by AlphaFold3 during complex prediction.

### 5.6. CryoEM Map analysis

For cryo-EM analysis, ProNA3D enables interface-focused analysis of density maps by zoning the map around the identified interface regions. This is implemented using the *volume zone* functionality of UCSF ChimeraX [29]. Cryo-EM maps are retrieved from the Electron Microscopy Data Bank (EMDB) and supplied to ProNA3D, along with the corresponding atomic models, allowing interface-based density zoning to be performed directly within the plug-in. For the calculation of the local resolution, we used RELION-4.0.0 [36] based on half maps.

### 5.7. Dataset curation for large-scale PDB analysis

PDB entries were initially filtered to structures containing nucleic acids with a resolution ≤ 3.0 Å, yielding 10,806 complexes. To reduce sequence redundancy, MMseqs2 [53] was employed in a multi-step protocol. Monomeric chains were first grouped by macromolecular type (protein, DNA, RNA). Protein sequences (derived from SEQRES records in the CIF file) were clustered at 30% sequence identity, whereas DNA and RNA sequences were clustered at 90% identity.

Subsequently, ProNA3D (default d_HA_ settings) was used to identify all potential interfaces within each complex (without interface clustering). Processing the dataset using the command-line implementation of ProNA3D required 5 h 51 min 51 s on a system equipped with an AMD EPYC 7552 48-Core Processor and 188 GB of RAM, running Ubuntu 24.04.4 LTS. Therefore, on average, a single complex takes less than 2 seconds to be processed on the same system. Interfaces were then mapped to cluster-cluster interactions, and one representative from each cluster combination was randomly selected, resulting in 6,642 complexes. In a final filtering step, sequence length thresholds were applied (≥ 30 residues for proteins and ≥ 6 nucleotides for nucleic acids), yielding a dataset of 6,047 complexes. Application of ProNA3D (default d_HA_ and d_C_ settings) to this dataset identified 59,959 sub-interfaces.

Finally, sub-interfaces were classified by interaction type as follows: RNA-RNA (n = 4,797), RNA-DNA (n = 450), DNA-DNA (n = 4,590), protein-DNA (n = 11,791), and protein-RNA (n = 23,616). Additionally, 14,681 interfaces were detected between protein chains only, and 34 sub-interfaces involved hybrid chains containing both amino acid residues and nucleotides that could not be unambiguously assigned to a single category (e.g., AA-tRNA). As ProNA3D was developed primarily for the analysis of nucleic acid-containing interfaces, these latter two interface classes were excluded, resulting in a final dataset of 45,244 sub-interfaces.

## 6. Acknowledgements

The authors thank the Topf group for helpful discussions and feedback. We especially thank Dr. Guendalina Marini and Dr. Mauro Maiorca for their support with the cryo-EM analysis.

## 9. Data availability

ProNA3D is implemented as a plug-in for the molecular visualization program UCSF ChimeraX and as a command-line tool. It is freely available for download from the ChimeraX toolshed and https://gitlab.com/topf-lab/ProNA3D. In addition, the repository contains the results of the large-scale analyses. Note that for secondary structure features, DSSR has to be downloaded with a license [21].

