## Supplementary material for "ProNA3D: Distance-Based Analysis of Nucleic Acid-Containing Interfaces": Suppl.

### Supplemental material

#### Supplemental figures

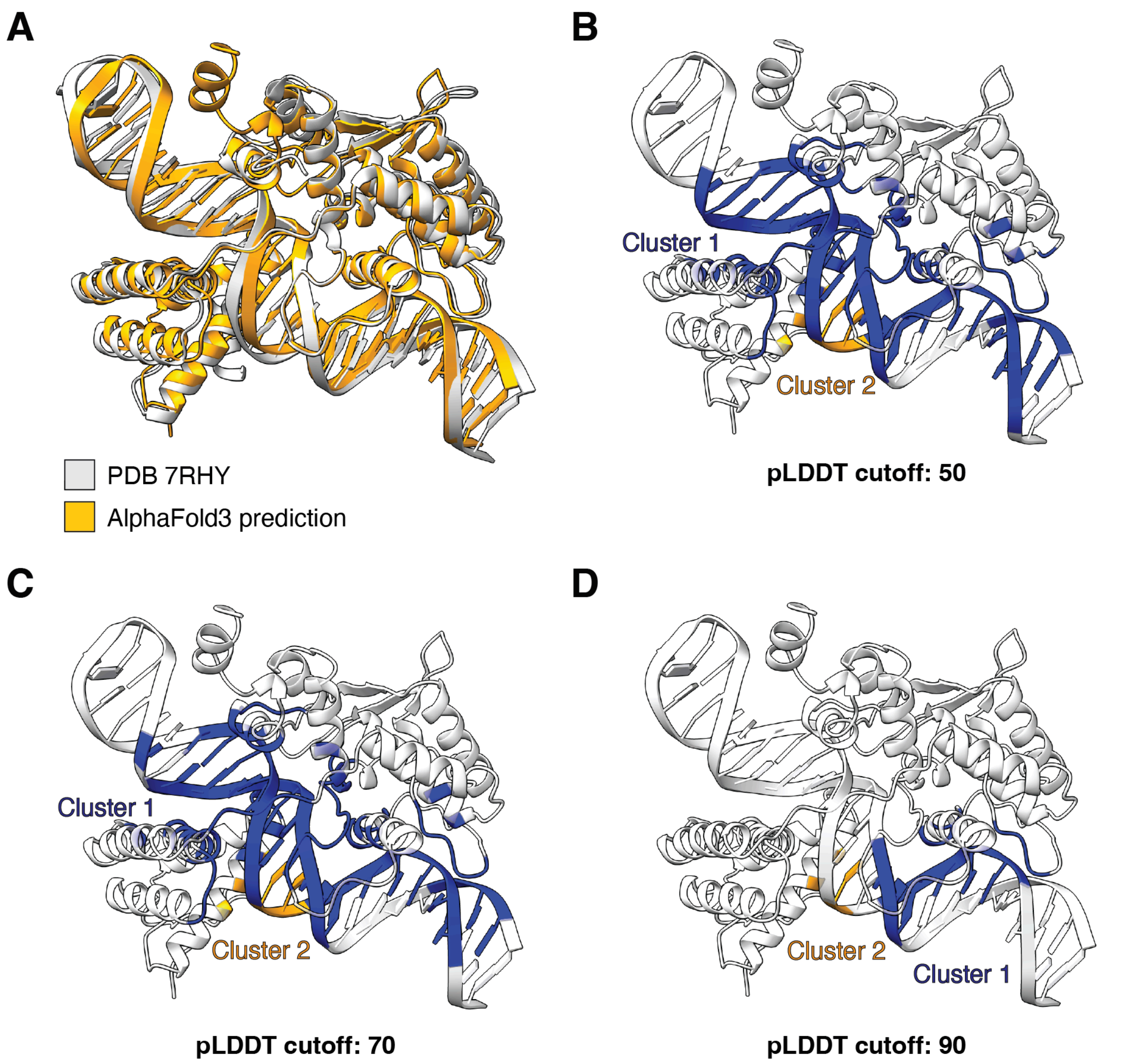


**Suppl. Figure 1:** ProNA3D analysis of the AlphaFold3-predicted complex of the Cre recombinase from *Punavirus P1* bound to its target DNA. **(A)** Superposition of the AlphaFold3 model (orange, Cre recombinase residues: 1-343, DNA nucleotides: 1-49) on the experimentally determined structure (PDB ID: 7RHY) (white, Cre recombinase residues: 20-329 with unresolved regions not modeled, DNA nucleotides: 1-49). **(B)** Sub-interfaces identified by ProNA3D using a pLDDT cutoff of 50, with sub-interface 1 (105 interacting residues and nucleotides) shown in blue and sub-interface 2 (5 interacting residues and nucleotides) in orange. **(C)** Sub-interfaces detected using a pLDDT cutoff of 70, colored as in (B) (Sub-interface 1: 95 interacting residues and nucleotides, Sub-interface 2: 5 interacting residues and nucleotides). **(D)** Sub-interfaces detected using a pLDDT cutoff of 90, with sub-interface coloring consistent with (B) (Sub-interface 1: 16 interacting residues and nucleotides, sub-interface 2: 3 interacting residues and nucleotides).

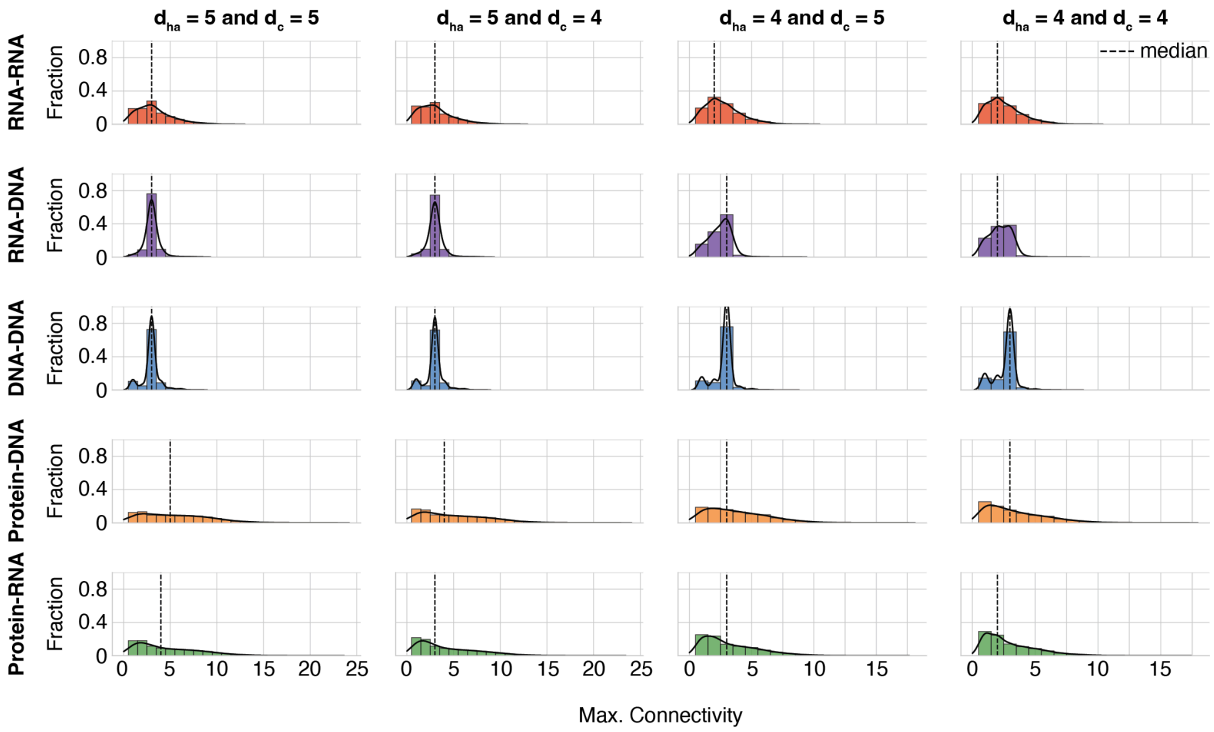


**Suppl. Figure 2:** Maximum connectivity across nucleic acid-containing interface types and varying $d_{HA}$ and $d_C$ thresholds. Connectivity was defined as the number of distinct residues or nucleotides contacted by a residue or nucleotide across an interface. For each interface, the highest connectivity value was extracted. Shown are RNA-RNA (red), RNA-DNA (purple), DNA-DNA (blue), protein-DNA (orange), and protein-RNA (green) interfaces. Median values are indicated for each interface class (dotted line). **(A)** $d_{HA}$ = 5 and $d_C$ = 5, **(B)** $d_{HA}$ = 5 and $d_C$ = 4, **(C)** $d_{HA}$ = 4 and $d_C$ = 5, **(D)** $d_{HA}$ = 4 and $d_C$ = 4.

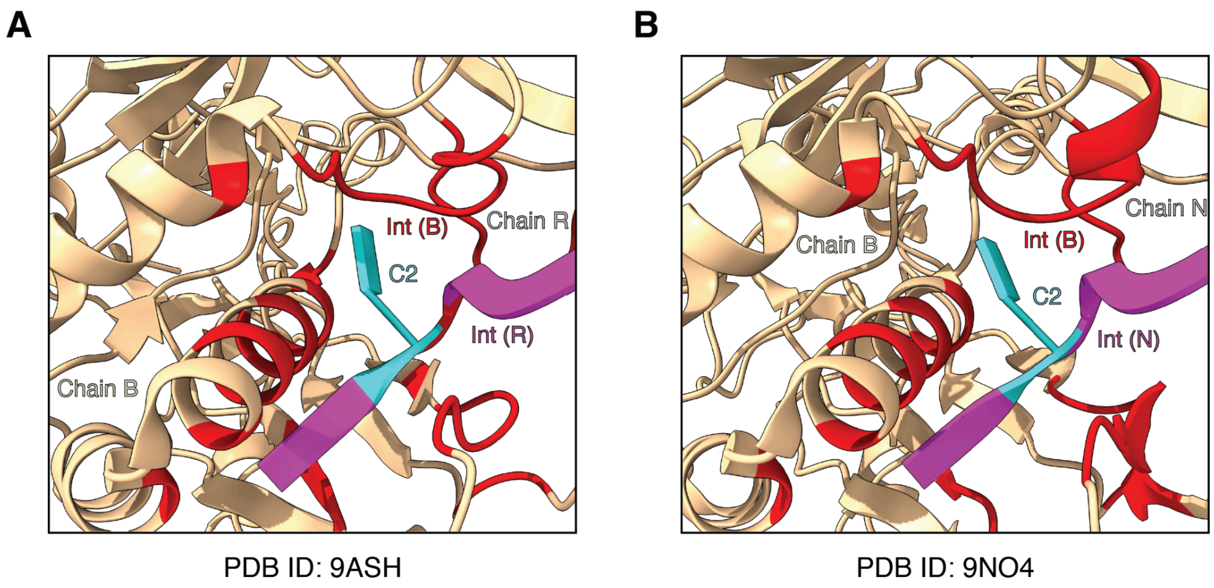


**Suppl. Figure 3:** Conserved high-connectivity nucleotide flipping in CRISPR-associated protein Csm4 complexes (red: Csm4 interface residues, magenta: RNA interface nucleotides, cyan: flipped high-connectivity nucleotide). **(A)** CRISPR-associated protein Csm4 from *L. lactis* bound to crRNA (PDB: 9ASH). **(B)** CRISPR-associated protein Csm4 from *S. thermophilus*bound to crRNA (PDB: 9NO4).

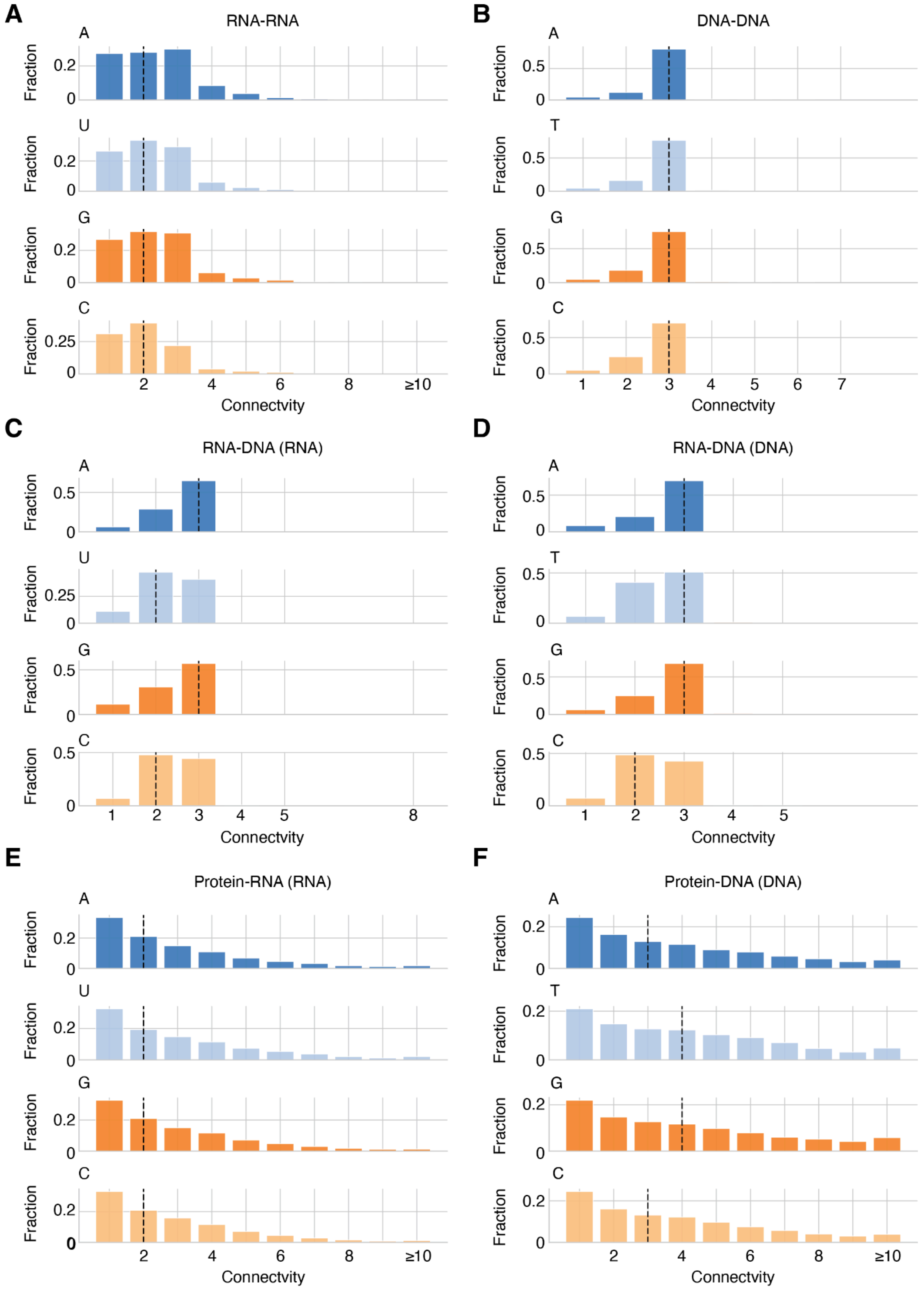


**Suppl. Figure 4:** Per-nucleotide type connectivity (A/DA: blue; U/DT: lightblue; DG/G: orange; C/DC: lightorange) across nucleic acid-containing interface types. Connectivity was defined as the number of distinct residues or nucleotides contacted by a residue or nucleotide across an interface. **(A)** RNA-RNA, **(B)** DNA-DNA, **(C)** RNA-DNA (RNA), **(D)** RNA-DNA (DNA), **(E)** protein-RNA (RNA), and **(F)** protein-DNA (DNA). Median values are indicated for each interface class (dotted line).

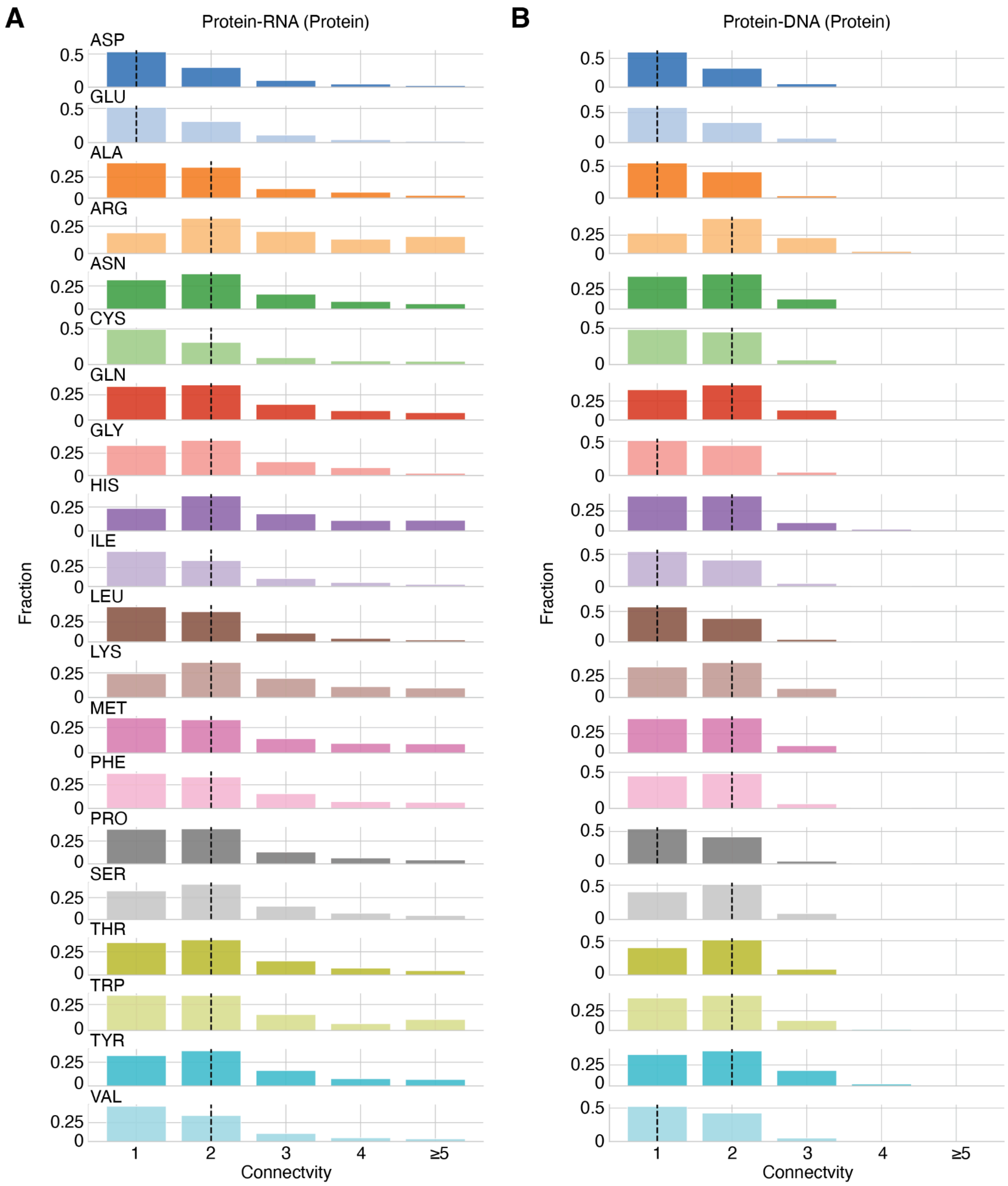


**Suppl. Figure 5:** Per-residue type connectivity across nucleic acid-containing interface types. Connectivity was defined as the number of distinct residues or nucleotides contacted by a residue or nucleotide across an interface. **(A)** protein-RNA (protein) and **(B)** protein-DNA (protein). Median values are indicated for each interface class (dotted line).

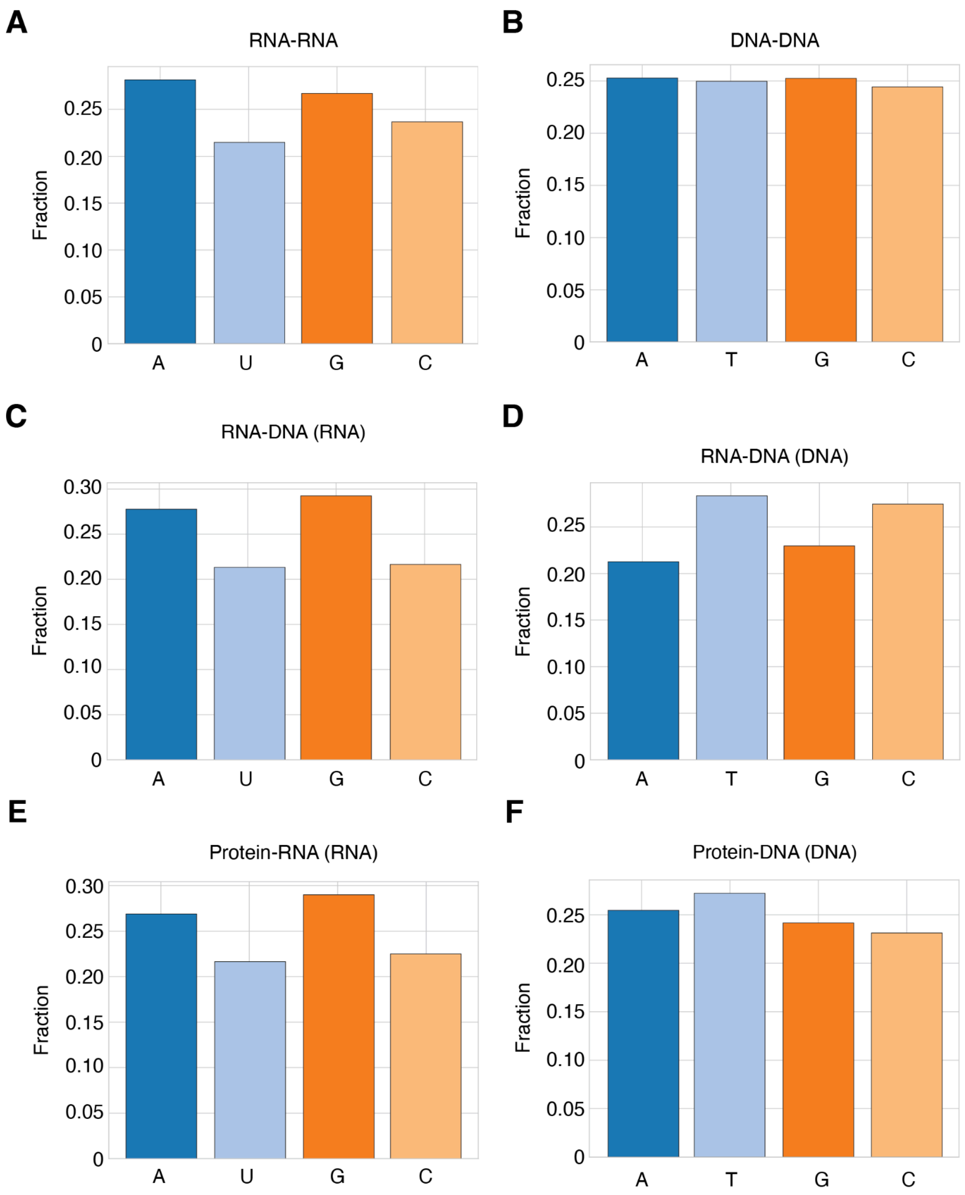


**Suppl. Figure 6:** Per-nucleotide type distributions (A: blue; U/T: lightblue; G: orange; C: lightorange) across nucleic acid-containing interface types. **(A)** RNA-RNA, **(B)** DNA-DNA, **(C)** RNA-DNA (RNA), **(D)** RNA-DNA (DNA), **(E)** protein-RNA (RNA), and **(F)** protein-DNA (DNA).

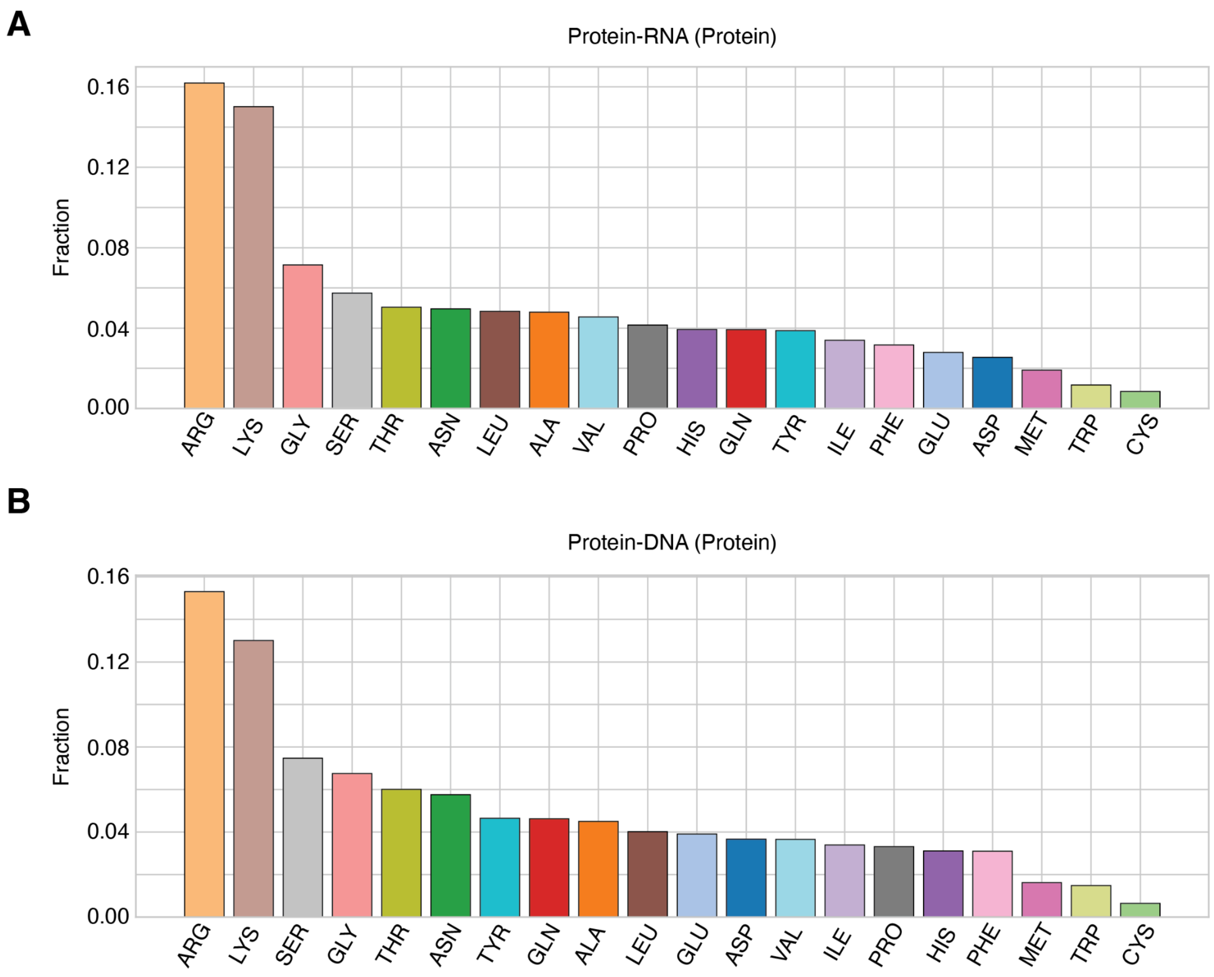


**Suppl. Figure 7:** Per-residue type distributions across nucleic acid-containing interface types. **(A)** protein-RNA (protein) and **(B)** protein-DNA (protein).

**Supplemental tables**

**Suppl. Table 1:** Comparison of available computational tools for structural analysis of nucleic acid-containing complexes.

| Tool name | Supported molecule types | Support for predicted structures (3D)* | Confidence metric integration | cryo-EM functionality | Topology visualization | Interactive visualization | command-line support |
|---|---|---|---|---|---|---|---|
| **DNAproDB** | Protein-DNA | Yes | No | No | Yes (for DNA) | Yes | Yes |
| **RNAproDB** | Protein-RNA, nucleic-acid-containing complexes | Yes | No | No | Yes (for nucleic acids) | Yes | Yes |
| **DSSR** | Nucleic acid-containing structures | Yes | No | No | Yes (main purpose) | Yes (wDSSR web interface) | Yes |
| **RNAscape** | Nucleic acid-containing structures | Yes | No | No | Yes | Yes | Yes |
| **ViennaRNA** | RNA sequences and RNA secondary structures | No | No | No | Yes | No (static plots only) | Yes |
| **PPI3D** | Protein-, peptide-, and nucleic | No | No | No | No | Yes | No |

| | | | | | | | |
|---|---|---|---|---|---|---|---|
| | acid-containing complexes | | | | | | |
| **PLIP** | Protein-, small-molecule-, nucleic acid-containing complexes | Yes | No | No | No | Yes | Yes |
| **VoroContacts** | Proteins, nucleic acids, and their complexes | Yes | No | No | No | No | Yes |
| **RNAView** | Nucleic acids | Yes | No | No | Yes | Yes | Yes |
| **ProNA3D** | Nucleic acid-containing complexes | Yes | Yes | Yes | Yes (DSSR-based) | Yes | Yes |

*Support for predicted structures refers to the ability to analyze user-provided predicted 3D coordinate models (e.g., AlphaFold3), rather than the generation of predicted structures or integration of prediction confidence scores.

**Suppl. Table 2:** Maximum connectivity across all interface types for $d_{HA}$ = 5 Å and $d_C$ = 5 Å. Shown are the number of sub-interfaces analysed (count) and the distribution of maximal connectivity values, reported as median, mean, and standard deviation.

| Interface type | Count | Median | Mean | Std |
|---|---|---|---|---|
| RNA-RNA | 4,797 | 3.00 | 3.22 | 1.83 |
| RNA-DNA | 450 | 3.00 | 3.03 | 0.76 |
| DNA-DNA | 4,590 | 3.00 | 2.92 | 0.88 |
| Protein-DNA | 11,791 | 5.00 | 5.52 | 3.48 |
| Protein-RNA | 23,616 | 4.00 | 4.64 | 3.32 |

**Suppl. Table 3:** Mann-Whitney U (Wilcoxon rank-sum) test for maximum connectivity interface type populations.

| P-value | RNA-RNA | DNA-DNA | RNA-DNA | Protein-DNA | Protein-RNA |
|---|---|---|---|---|---|
| **RNA-RNA** | x | 1.18e-01 | 3.84e-01 | << 0.001 | 8.49e-110 |
| **DNA-DNA** | x | x | 1.04e-01 | << 0.001 | 1.87e-114 |
| **RNA-DNA** | x | x | x | 3.74e-46 | 1.83e-10 |
| **Protein-DNA** | x | x | x | x | 3.77e-125 |
| **Protein-RNA** | x | x | x | x | x |

**Suppl. Table 4:** Maximum connectivity across all interface types for $d_{HA}$ = 5 Å and $d_C$ = 4 Å. Shown are the number of sub-interfaces analysed (count) and the distribution of maximal connectivity values, reported as median, mean, and standard deviation.

| Interface type | Count | Median | Mean | Std |
|---|---|---|---|---|
| RNA-RNA | 5,469 | 3.00 | 3.06 | 1.81 |
| RNA-DNA | 460 | 3.00 | 3.00 | 0.78 |
| DNA-DNA | 4,685 | 3.00 | 2.91 | 0.89 |

| | | | | |
|---|---|---|---|---|
| Protein-DNA | 13,792 | 4.00 | 5.06 | 3.47 |
| Protein-RNA | 29,264 | 3.00 | 4.26 | 3.21 |

**Suppl. Table 5:** Maximum connectivity across all interface types for $d_{HA}$ = 4 Å and $d_C$ = 5 Å. Shown are the number of sub-interfaces analysed (count) and the distribution of maximal connectivity values, reported as median, mean, and standard deviation.

| Interface type | Count | Median | Mean | Std |
|---|---|---|---|---|
| RNA-RNA | 4,954 | 2.00 | 2.67 | 1.35 |
| RNA-DNA | 546 | 3.00 | 2.43 | 0.84 |
| DNA-DNA | 4,915 | 3.00 | 2.76 | 0.75 |
| Protein-DNA | 13,555 | 3.00 | 3.80 | 2.38 |
| Protein-RNA | 31,894 | 3.00 | 3.26 | 2.22 |

**Suppl. Table 6:** Maximum connectivity across all interface types for $d_{HA}$ = 4 Å and $d_C$ = 4 Å. Shown are the number of sub-interfaces analysed (count) and the distribution of maximal connectivity values, reported as median, mean, and standard deviation.

| Interface type | Count | Median | Mean | Std |
|---|---|---|---|---|
| RNA-RNA | 6,088 | 2.00 | 2.51 | 1.34 |
| RNA-DNA | 781 | 2.00 | 2.21 | 0.85 |
| DNA-DNA | 5,832 | 3.00 | 2.64 | 0.81 |
| Protein-DNA | 17,375 | 3.00 | 3.39 | 2.31 |
| Protein-RNA | 45,631 | 2.00 | 2.97 | 2.07 |

**Suppl. Table 7:** Connectivity statistics by nucleotide type for DNA-DNA interfaces.

| Interface type | Count | Median | Mean | Std |
|---|---|---|---|---|
| A | 28,518 | 3.00 | 2.82 | 0.60 |

| | | | | |
|---|---|---|---|---|
| T | 28,153 | 3.00 | 2.78 | 0.61 |
| G | 28,473 | 3.00 | 2.75 | 0.60 |
| C | 27,562 | 3.00 | 2.68 | 0.61 |

**Suppl. Table 8:** Connectivity statistics by nucleotide type for RNA-RNA interfaces.

| Interface type | Count | Median | Mean | Std |
|---|---|---|---|---|
| A | 15,874 | 2.00 | 2.44 | 1.31 |
| U | 12,110 | 2.00 | 2.31 | 1.17 |
| G | 15,056 | 2.00 | 2.34 | 1.17 |
| C | 13,342 | 2.00 | 2.13 | 1.10 |

**Suppl. Table 9:** Connectivity statistics by nucleotide type (RNA) for RNA-DNA interfaces.

| Interface type | Count | Median | Mean | Std |
|---|---|---|---|---|
| A | 1,364 | 3.00 | 2.60 | 0.63 |
| U | 1,047 | 2.00 | 2.31 | 0.68 |
| G | 1,436 | 3.00 | 2.47 | 0.72 |
| C | 1,063 | 2.00 | 2.39 | 0.63 |

**Suppl. Table 10:** Connectivity statistics by nucleotide type (DNA) for RNA-DNA interfaces.

| Interface type | Count | Median | Mean | Std |
|---|---|---|---|---|
| A | 1,014 | 3.00 | 2.64 | 0.66 |
| T | 1,351 | 3.00 | 2.49 | 0.70 |
| G | 1,095 | 3.00 | 2.64 | 0.62 |
| C | 1,309 | 2.00 | 2.39 | 0.66 |

**Suppl. Table 11:** Connectivity statistics by nucleotide type (DNA) for protein-DNA interfaces.

| Interface type | Count | Median | Mean | Std |
|---|---|---|---|---|

| | | | | |
|---|---|---|---|---|
| A | 14,097 | 3.00 | 3.88 | 2.78 |
| T | 15,072 | 4.00 | 4.15 | 2.81 |
| G | 13,374 | 4.00 | 4.20 | 2.96 |
| C | 12805 | 3.00 | 3.84 | 2.74 |

**Suppl. Table 12:** Connectivity statistics by residue type (protein) for protein-DNA interfaces.

| Interface type | Count | Median | Mean | Std |
|---|---|---|---|---|
| ASP | 4,807 | 1.00 | 1.46 | 0.63 |
| GLU | 5,114 | 1.00 | 1.51 | 0.68 |
| ALA | 5,898 | 1.00 | 1.49 | 0.59 |
| ARG | 20,052 | 2.00 | 2.02 | 0.82 |
| ASN | 7,539 | 2.00 | 1.73 | 0.72 |
| CYS | 845 | 2.00 | 1.58 | 0.62 |
| GLN | 6,056 | 2.00 | 1.75 | 0.72 |
| GLY | 8,843 | 1.00 | 1.55 | 0.60 |
| HIS | 4,070 | 2.00 | 1.72 | 0.76 |
| ILE | 4,443 | 1.00 | 1.51 | 0.60 |
| LEU | 5,264 | 1.00 | 1.47 | 0.59 |
| LYS | 17,042 | 2.00 | 1.74 | 0.72 |
| MET | 2,123 | 2.00 | 1.66 | 0.68 |
| PHE | 4,061 | 2.00 | 1.63 | 0.63 |
| PRO | 4,337 | 1.00 | 1.51 | 0.59 |
| SER | 9,791 | 2.00 | 1.69 | 0.64 |
| THR | 7,872 | 2.00 | 1.70 | 0.65 |
| TRP | 1,945 | 2.00 | 1.74 | 0.73 |
| TYR | 6,092 | 2.00 | 1.86 | 0.80 |

| | | | | |
|---|---|---|---|---|
| VAL | 4,789 | 1.00 | 1.53 | 0.60 |

**Suppl. Table 13:** Connectivity statistics by nucleotide type (RNA) for protein-RNA interfaces.

| Interface type | Count | Median | Mean | Std |
|---|---|---|---|---|
| A | 68,085 | 2.00 | 2.96 | 2.26 |
| U | 54,827 | 2.00 | 3.09 | 2.35 |
| G | 73,444 | 2.00 | 2.93 | 2.16 |
| C | 56,965 | 2.00 | 2.89 | 2.11 |

**Suppl. Table 14:** Connectivity statistics by residue type (protein) for protein-RNA interfaces.

| Interface type | Count | Median | Mean | Std |
|---|---|---|---|---|
| ASP | 8,397 | 1.00 | 1.75 | 1.04 |
| GLU | 9,224 | 1.00 | 1.76 | 1.01 |
| ALA | 15,824 | 2.00 | 1.95 | 1.12 |
| ARG | 53,411 | 2.00 | 2.87 | 1.62 |
| ASN | 16,357 | 2.00 | 2.21 | 1.21 |
| CYS | 2,806 | 2.00 | 1.87 | 1.15 |
| GLN | 12,961 | 2.00 | 2.27 | 1.32 |
| GLY | 23,561 | 2.00 | 2.08 | 1.07 |
| HIS | 12,987 | 2.00 | 2.58 | 1.48 |
| ILE | 11,211 | 2.00 | 1.86 | 1.09 |
| LEU | 15,944 | 2.00 | 1.85 | 1.01 |
| LYS | 49,536 | 2.00 | 2.53 | 1.41 |
| MET | 6,315 | 2.00 | 2.33 | 1.48 |
| PHE | 10,426 | 2.00 | 2.17 | 1.30 |
| PRO | 13,701 | 2.00 | 2.05 | 1.19 |

| | | | | |
|---|---|---|---|---|
| SER | 18,946 | 2.00 | 2.12 | 1.14 |
| THR | 16,630 | 2.00 | 2.11 | 1.17 |
| TRP | 3,878 | 2.00 | 2.38 | 1.61 |
| TYR | 12,774 | 2.00 | 2.24 | 1.30 |
| VAL | 15,037 | 2.00 | 1.86 | 1.10 |
